# Three small partner proteins facilitate the type VII-dependent secretion export of an antibacterial nuclease

**DOI:** 10.1101/2023.04.01.535202

**Authors:** Yaping Yang, Eleanor Boardman, Justin Deme, Felicity Alcock, Susan Lea, Tracy Palmer

## Abstract

The type VIIb protein secretion system (T7SSb) plays a role in interbacterial competition in Gram-positive Firmicute bacteria and secretes various toxic effector proteins. The mechanism of secretion and the roles of numerous conserved genes within T7SSb gene clusters remain unknown. EsaD is a nuclease toxin secreted by the *Staphylococcus aureus* T7SSb, which forms a complex with its cognate immunity protein, EsaG, and chaperone EsaE. Encoded upstream of EsaD are three small secreted proteins, EsxB, EsxC and EsxD. Here we show that EsxBCD bind to the transport domain of EsaD and function as EsaD export factors. We report the first structural information for a complete T7SSb substrate pre-secretion complex. Cryo-EM of the EsaDEG trimer and the EsaDEG-EsxBCD hexamer shows that incorporation of EsxBCD confers a conformation comprising a flexible globular cargo domain attached to a long narrow shaft that is likely to be crucial for efficient toxin export.

## Introduction

In mixed bacterial communities interbacterial antagonism plays a central role in determining the composition of the microbiota.^1^ Bacteria have evolved a diverse array of toxins, secretion systems, and defense mechanisms, for survival in what can be highly competitive environments.^2^ The T7SS is a protein secretion system that is frequently found in Gram-positive bacteria and is classified into two distantly related subtypes.^3^ The T7SSa is found across actinobacterial species and was first described in pathogenic mycobacteria where it is essential for virulence.^4–6^ The T7SSb system is found in firmicutes, in both pathogenic (e.g. *Staphylococcus aureus*) and non-pathogenic (e.g. *Bacillus subtilis*) species.^7, 8^ The primary role of the T7SSb is inhibition of competitor bacteria via secretion of antibacterial toxins.^9–15^ These toxin substrates usually have a transport-associated N-terminal LXG domain (IPR006829), and a highly polymorphic C-terminal toxin domain. T7b-associated toxin domains identified to date include nucleases, membrane depolarizing toxins, lipid II phosphatases, NADases and ADP ribosyltransferases.^9–11,13^

The *S. aureus* T7SSb is encoded at the *ess* locus.^8^ The secretion system components are encoded at the 5’ end of the gene cluster and are conserved across T7^+^ *S. aureus* strains. The central component of the machinery is EssC, a membrane-bound AAA+ ATPase of the SpoIIIE/FtsK family. Based on similarity to the EccC component of the T7SSa, an EssC hexamer likely forms the T7b transport channel.^16, 17^ The C-terminal region of EssC shows significant diversity across *S. aureus* strains and can classified into one of four types, EssC1 to EssC4.^18^ The first gene in the *ess* locus is *esxA*, encoding a WXG100 protein (IPR036689). The WXG100 family is associated with T7-secretion, and EsxA is critical to the function of the core machinery in both the T7SSa and T7SSb. While EsxA is essential for substrate export, it is itself secreted by the T7 apparatus.^8, 19^ The 3’ region of the *ess* locus is more variable, with sequences diverging at the 3’ end of the *essC* gene (Fig. 1A). Variant-specific genes found downstream of *essC* include secreted substrates, immunity proteins, and a number of genes of as yet unknown function.

**Figure 1.**
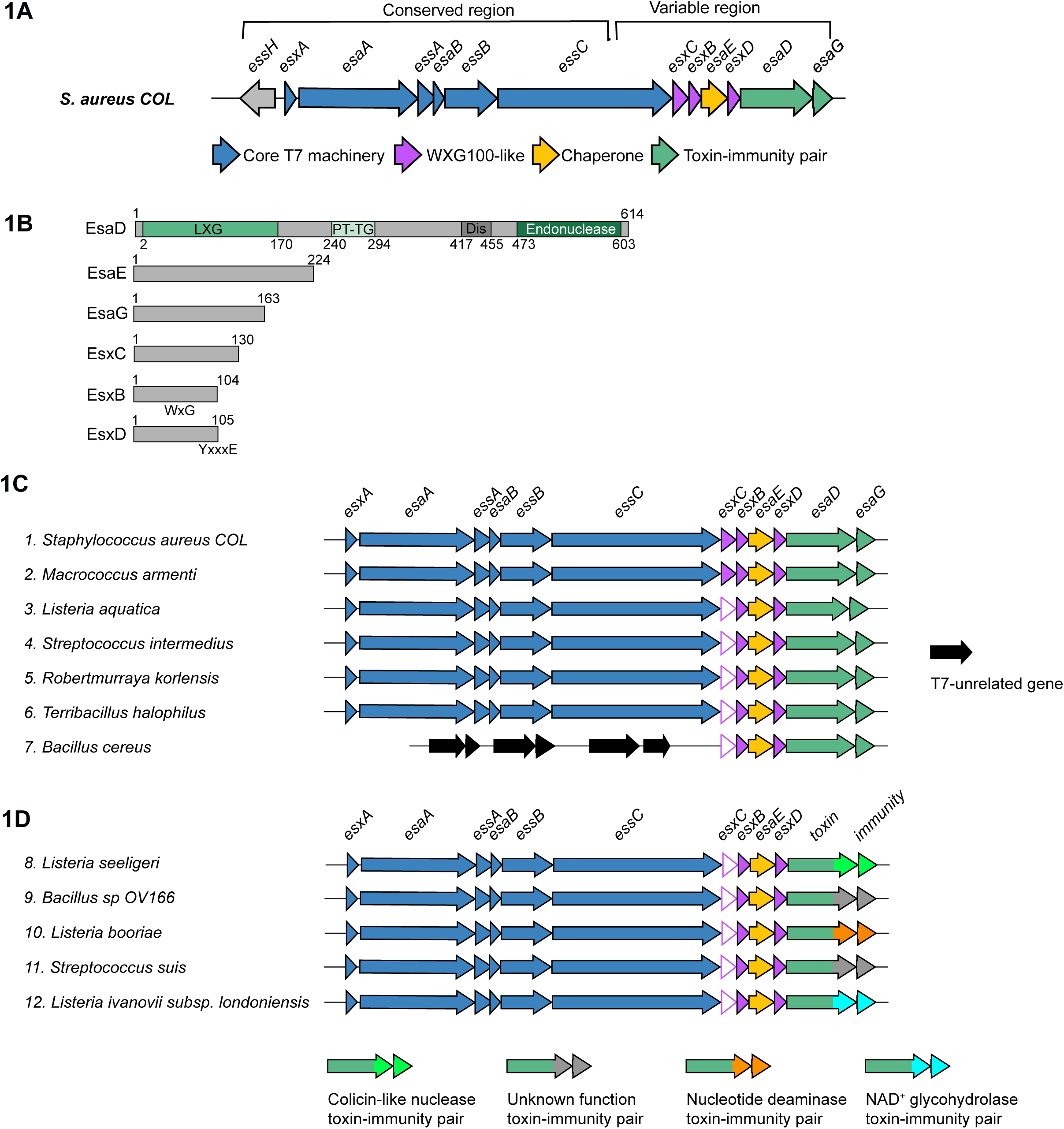
*esxB*, *esxC* and *esxD* co-occur with *esaD*. A. Schematic of the T7 locus in the *essC-1* strain COL. The 5’ region from *essH* to approximately two thirds of the way into *essC* is highly conserved across all strains. The 3’ end of the locus encodes different variant-specific proteins, which for *essC-1* strains includes the *esaD-esaG* toxin-immunity pair. B. Schematic representation of *essC-1* specific proteins. The locations of EsaD LXG, PT-TG, disordered (‘Dis’) and endonuclease domains are indicated. The approximate positions of the EsxB WXG motif and EsxD targeting signal (YxxxE) are indicated. C and D. Representative loci from gene neighborhood analysis of *esaD*. Genes corresponding to *esxB*, *esxC* and *esxD* occur in a conserved four-gene cluster (*esxC-esxB-esaE-esxD*) directly upstream of either *esaD* (C), or an *esaD*-like gene with a different toxin domain (D). N.B. *esxC* genes were designated as two distinct groups by WebFlaGs (closed/open pink arrows). However, the conserved gene size, synteny and predicted fold (Fig. S1A) all indicate that both groups are *esxC*. Accessions for the EsaD-like protein from each cluster are as follows: (1) WP_000159025, (2) WP_224186791, (3) WP_244964453, (4) WP_112384664, (5) WP_251523463, (6) WP_077309997, (7) WP_151522567, (8) WP_194333108, (9) WP_088085129, (10) WP_185646797, (11) WP_256797855, (12) WP_077916384.

EsaD is an antibacterial nuclease found in *essC1* strains of *S. aureus* and is related to the *B. subtilis* YeeF toxin.^9^ It has helical N-terminal region, a central PT-TG domain (IPR027797) and a C-terminal nuclease domain (IPR044927) (Fig. 1B). EsaG, the EsaD immunity protein is encoded downstream of *esaD*. Sandwiched between *essC1* and *esaD* are four genes: *esxC*, *esxB*, *esaE* and *esxD* (Fig. 1A). EsaE is a chaperone which binds to EsaD and targets it to the T7 apparatus^9^, and EsxB is small α-helical hairpin of the WXG100 family. EsxC and EsxD, while not annotated as WXG100 proteins, are similarly predicted to fold as small α-helical hairpins^20^, and EsxD has a putative T7SS targeting signal (YxxxD/E) at its C-terminus (Fig. 1B).^21, 22^ Prior work has shown that EsxB, EsxC and EsxD are all secreted by the *S. aureus* T7SS and both EsxB and EsxC have been implicated as effectors involved in the pathogenesis of *S. aureus* murine abscesses.^8, 20, 23^

Here we have further investigated the functions of the small WXG100-like Esx proteins EsxB, EsxC and EsxD. We show that they form a complex with EsaDEG, and that they stabilize and dramatically boost EsaD secretion. To gain insight into the secretion mechanism we undertook structural analysis of EsaDEG and EsaDEG-EsxBCD complexes using cryo-electron microscopy. Although the flexible nature of the complexes precluded high-resolution structure determination, the low-resolution maps show that incorporation of EsxBCD gives the complex a distinctive long, narrow conformation, similar to the PE/PPE substrates of the mycobacterial T7SSa.^24^ We propose that this is a common structural feature of T7-secreted substrates from both T7a and T7b systems.

## Results

### EsxB, EsxC and EsxD are export/stability factors for EsaD

Prior work has implicated the small, secreted substrates, EsxB and EsxC in *S. aureus* virulence.^8, 23^. EsxD has also been identified as a T7SS substrate, and an interaction partner of EsxB.^20^ Gene neighborhood analysis reveals that genes encoding homologs of these three small WXG100-like proteins always co-occur with *esaD*, and with conserved synteny (*esxC-esxB-esaE-esxD*). While the gene cluster is frequently found at the T7SS locus in firmicutes, in some *Bacillus* species the cluster is found elsewhere on the genome (Fig. 1C). Of the three small proteins, EsxC is the most divergent in sequence, although structural predictions indicate a conserved fold (Fig. S1).

As EsxB, EsxC and EsxD are also encoded by non-pathogenic bacteria such as *Listeria aquatica*, they are unlikely to have a conserved role in pathogenesis. Instead, based on the co-occurrence with EsaD and EsaE, we hypothesized that they might function in secretion of EsaD by the T7SS. We have previously shown that EsaD is secreted, albeit inefficiently, when EsaD, EsaE and EsaG are overproduced from a plasmid in *S. aureus essC1* strains.^9^ To test our hypothesis, we constructed a plasmid carrying the whole genomic region from *esxC* to *esaG* under the control of the P(*xyl*/*tet*) promoter (pCBED-DG) (Fig. 2A). The *esaD* gene in this construct was modified to encode a C-terminal HA tag for immunodetection and carried the catalytic mutation H528A to reduce toxicity. We then made four additional constructs, each with a single gene deletion of *esxB*, *esxC*, *esxD* or *esaE.* These plasmids were transferred into the *S. aureus essC1* strain COL, in which the same genomic region (*esxC-esaG*) had been deleted (COLΔ*esxC-esxG*). Following induction of gene expression, the whole cell and culture supernatant fractions were isolated and analyzed for the presence of EsaD^H528A^HA. Immunoblot of these fractions showed that in the presence of all four accessory proteins, EsaD^H528A^HA was readily detected in both cell and supernatant fractions indicating secretion of EsaD^H528A^HA (Fig. 2B). Deletion of either *esxB* or *esxC* from the plasmid vastly reduced the EsaD^H528A^HA detected in the secreted fraction. Deletion of *esxD* or *esxE* produced an even more drastic effect, with EsaD^H528A^HA not detected (*ΔesxD*) or barely detected (*ΔesaE*) in the culture supernatant. In the case of all four deletions, the EsaD^H528A^HA signal in the cell fractions was also significantly reduced - in the case of Δ*esxD* to below the detection limit in this particular experiment, though a longer exposure showed that EsaD^H528A^HA was still minimally produced and secreted in this sample (Fig. 2B). A control immunoblot with antibodies against EsxA confirmed that the T7SS secretion efficiency was unaffected by the presence or absence of EsxB, EsxC and EsxD (Fig. 2B, middle panel). A similar approach showed that the addition of *esxB, esxC* and *esxD* to a plasmid carrying *esaE*, *esaD* and *esaG* vastly improves the secretion efficiency of EsaD (Fig. S2). These results clearly demonstrate that EsxB, EsxC, EsxD and EsaE all contribute to both the stability and secretion of EsaD, but not EsxA, and so can be considered EsaD-specific export factors.

**Figure 2.**
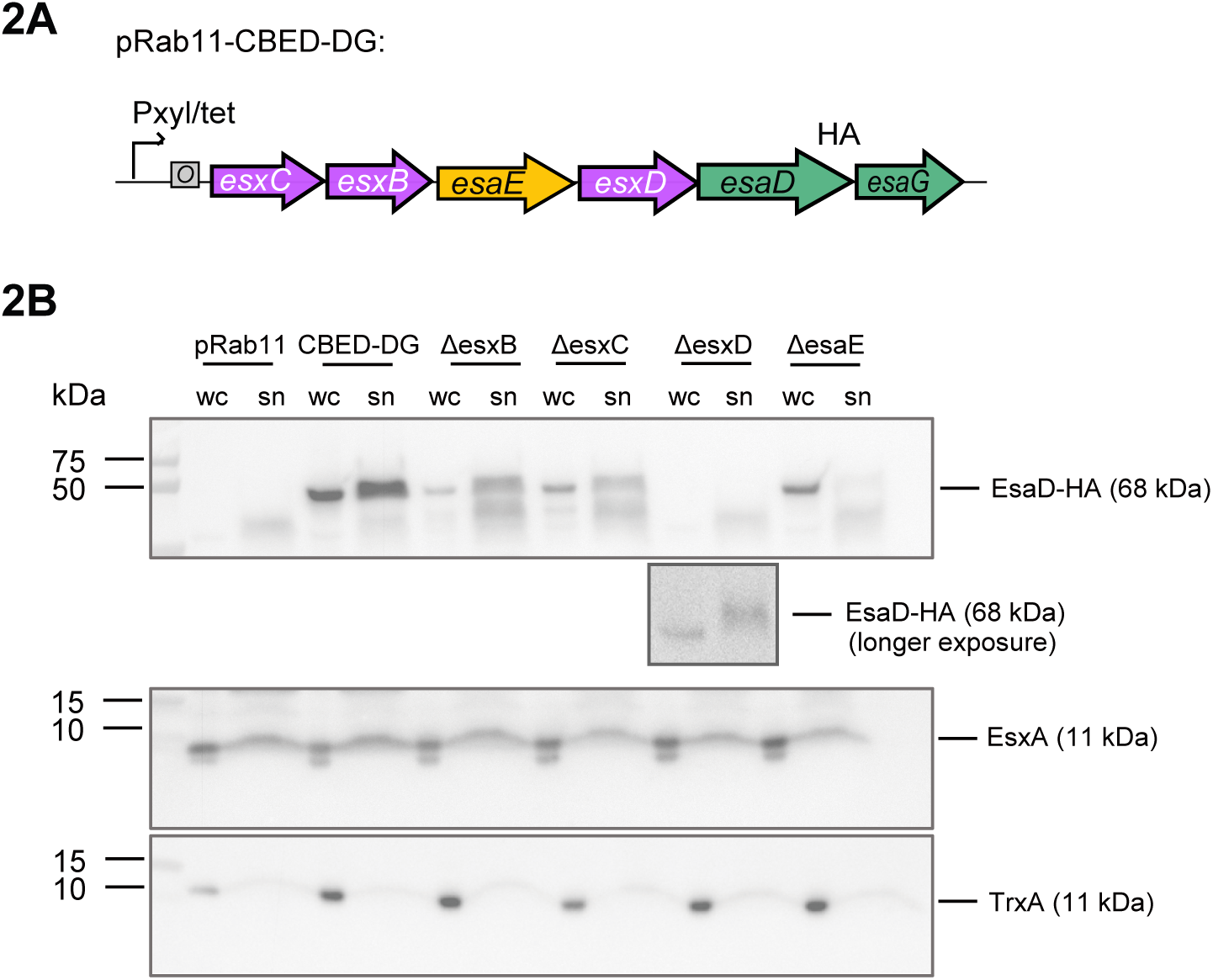
Contribution of EsxB, EsxC, EsaE and EsxD to EsaD secretion. A. Schematic representation of the coding region of plasmid pRab11-CBED-DG used in (B). ‘HA’ denotes a C-terminal HA epitope tag on EsaD^H528A^ B. COLΔ*esxC-esaG* cells carrying empty vector (pRab11), the complete pRab11-CBED-DG plasmid shown in (C) (CBED-DG) or derivatives of that plasmid with one of *esxB*, *esxC*, *esxD* or *esaE* deleted (*ΔesxB*, *ΔesxC*, *ΔesxD*, *ΔesaE*, respectively) were cultured in TSB growth medium. Following induction of plasmid-encoded gene expression, whole cell (wc) and culture supernatant (sn) fractions were isolated, prepared for immunoblot, and separated by SDS PAGE. EsaD was detected with antibodies against the C-terminal HA tag (top panel). A longer exposure of the anti-HA blot for the *ΔesxD* sample is shown below this panel. The T7-secreted control EsxA and cytoplasmic control TrxA were detected with anti-EsxA and anti-TrxA antibodies (middle and bottom panels). N.B. the supernatant fractions analyzed here correspond to 6 x more culture volume than the cell fractions.

### Composition of the EsaD pre-secretion complex

We have previously shown that prior to secretion, EsaD forms a ternary complex with its immunity protein EsaG and targeting chaperone EsaE.^9^ Given our observation that EsxB, EsxC and EsxD facilitate EsaD secretion and/or stability, and the fact that EsxB, EsxC and EsxD are themselves secreted by the T7SS^8, 11, 23, 25^, we explored whether the WXG100-like proteins could interact with the EsaDEG complex. All six genes were assembled on an *E. coli* expression plasmid where EsaD^H528A^ is encoded with a C-terminal his_6_ tag, and EsxD with a C-terminal twin-strep tag (Fig. 3A). Protein complexes were purified sequentially via the his-tag on EsaD^H528A^ and the twin-strep tag on EsxD and were analyzed by size exclusion chromatography (SEC) (Fig. 3B). A complex was eluted at a size corresponding to ∼400 kDa, significantly larger than the predicted molecular weight of a hetero-hexamer containing one copy of each protein (∼150 kDa). This complex was visualized by Coomassie staining (Fig. 3C) and all six components were identified by immunoblot (EsaD, EsxC, EsxD and EsxB) (Fig. 3D) or mass spectrometry (EsaE, EsaG).

**Figure 3.**
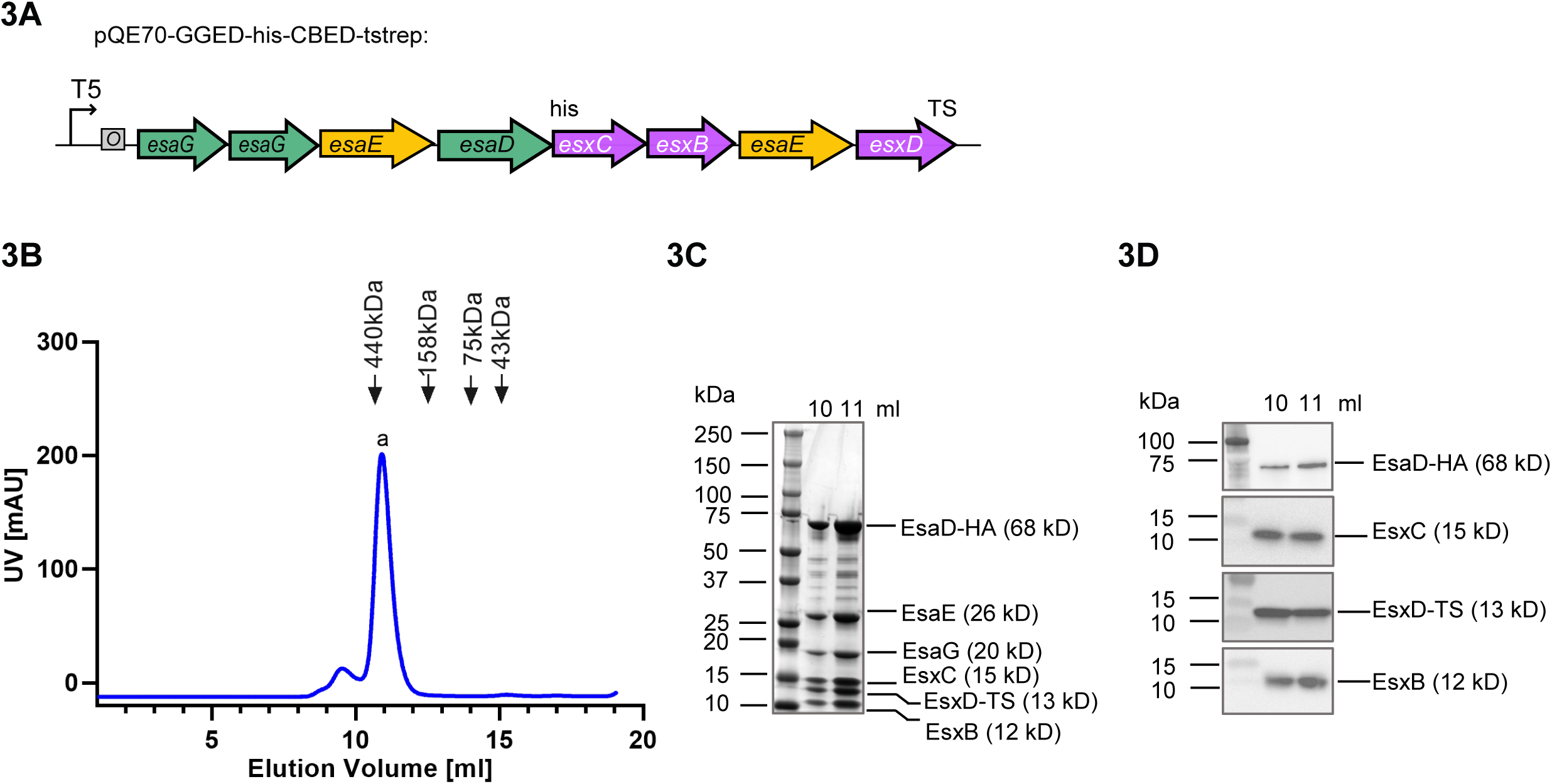
Purification of an EsaD hetero-hexamer. B. Schematic representation of the pQE70-based plasmid used for co-expression of *esxB, esxC, esxD, esaD, esaE* and *esaG*. ‘6xHis’ denotes a C-terminal His_6_ affinity tag on EsaD^H528A^ and ‘TS’ denotes a C-terminal twin-strep affinity tag on EsxD. C. *E. coli* M15 pREP4 cells carrying pQE70-GGEDCBED (shown in (A)) were grown in LB medium. Following induction of protein expression with IPTG, complexes were sequentially purified by nickel- and streptactin-affinity purification, and the resulting sample was analyzed by size-exclusion chromatography (SEC). D. Fractions corresponding to the major SEC peak at 10-12 ml in (A) were separated by SDS PAGE and visualized by Coomassie-staining. EsaG and EsaE were identified by mass spectrometry. E. Fractions corresponding to the major SEC peak at 10-12 ml in (A) were analyzed by antibodies against the his-tag (for EsaD-his), EsxC, the strep-tag (for EsxD-TS) and EsxB.

Previously it was noted that other toxin families with homology to *B. subtilis* YeeF / *S. aureus* EsaD were encoded in *Listeria monocytogenes* strains^13^, and Fig. 1D shows that they are also found in other firmicute species. These toxin sequences diverge prior to the nuclease domain, instead harboring other C-terminal toxin domains such as nucleotide deaminase or NAD^+^ glycohydrolase domains. Despite this, each of these toxin families is also encoded at a locus with *esxC-esxB-esaE-esxD* genes (Fig. 1D).^13^ We therefore reasoned that these accessory proteins may interact with the conserved N-terminal region of EsaD, rather than the nuclease domain. To address this, we repeated the co-expression of all six genes but with a truncated *esaD* lacking the C-terminal toxin sequence. As shown in Fig. 4A, EsxB, EsxC, EsxD and EsaE still co-purified with EsaD_1-477_-his by nickel affinity chromatography, indicating that these four partners interact with the transport-associated N-terminal domain of EsaD. When the N-terminal part of EsaD was removed, the EsaG immunity protein remained bound to the EsaD_421-614_-his nuclease as expected, but all other partners were lost (Fig. 4B).

**Figure 4.**
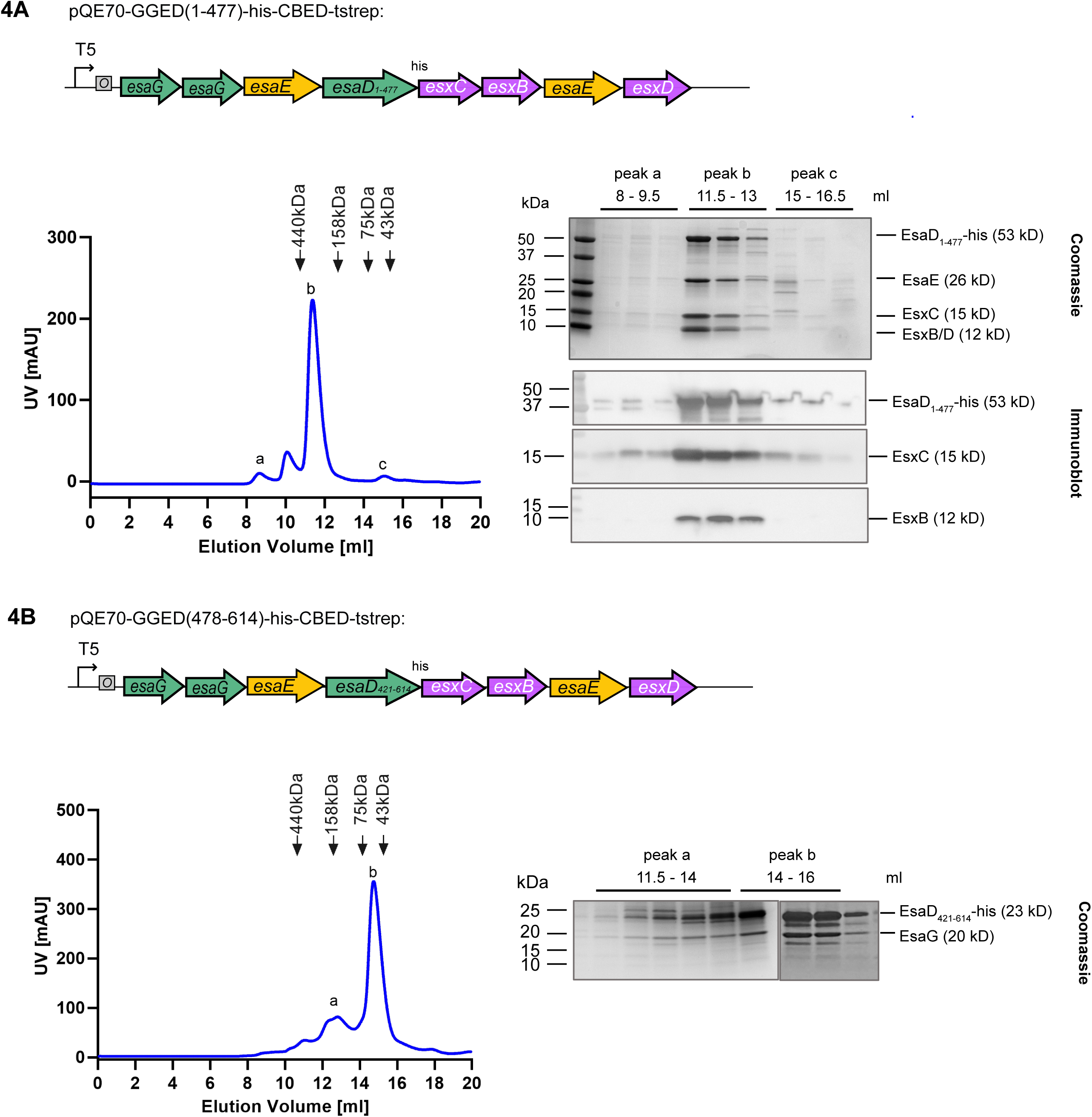
EsxB, EsxC, EsxD and EsaE co-purify with the EsaD N-terminal domain. A and B. *E. coli* M15 pREP4 cells carrying pQE70-GGED(1-477)-his-CBED (A) or pQE70-GGED(478-614)-his-CBED (B) were grown in LB medium, and protein expression was induced with IPTG. Complexes were purified by nickel affinity chromatography and analyzed by SEC (left hand panels). Fractions corresponding to the indicated elution volumes were analyzed by SDS-PAGE and Coomassie staining (right hand side, upper panels), or by immunoblot with antibodies against the his-tag (for EsaD-his), EsxC or EsxB ((A), right hand side, lower panel). EsxD in (A) and EsaG in (B) were identified by mass spectrometry.

### Structural analysis of the EsaD pre-secretion complex

To gain further insight into the EsaD secretion mechanism we analyzed the structures of EsaD-containing complexes. EsaD alone is unstable to purification, however in the presence of EsaE and EsaG a trimeric complex could be isolated (Fig. 5A). Most of the purified protein was eluted from the SEC column as a major peak at a volume corresponding to ∼400 kDa, much larger than the theoretical molecular weight of ∼100 kDa for a 1:1:1 heterotrimer (Fig. 5A, peak b, black trace). An additional smaller peak was eluted at an even larger molecular weight (Fig. 5A, peak a, black trace). Initial cryo-EM analysis of the ∼400 kDa peak fractions for yielded poor, undefined 2D class averages due to sample heterogeneity (not shown). Therefore, the EsaDEG trimer and the hexameric EsxBCD-EsaDEG complex were each subject to crosslinking with glutaraldehyde to reduce flexibility and improve particle alignment (Fig. 5A, B). Crosslinking of EsaDEG resulted in a later elution volume of EsaDEG, more consistent with the predicted molecular weight, and SDS-PAGE revealed the presence of ∼100 kDa EsaDEG trimers (Fig. 5A, peak b) and ∼200 kDa dimers-of-trimers (Fig 5A, peak a). Some additional higher molecular weight species were also present in the earliest fractions. Fractions corresponding to crosslinked EsaDEG trimers were analyzed by cryo-EM, discussed below. Glutaraldehyde crosslinking of the EsaDEG-EsxBCD hexamer (Fig 5B) produced two species which correspond in size to the heterohexamer (approximately 150 kDa) and dimers-of-hexamers (approximately 300 kDa), as judged by SDS-PAGE. Larger species might also be present but were not resolved on this gel (Fig 5B, right hand panel Crosslinked fractions corresponding to the ∼150 kDa hexamer (peak c) were selected for analysis by cryo-EM.

**Figure 5.**
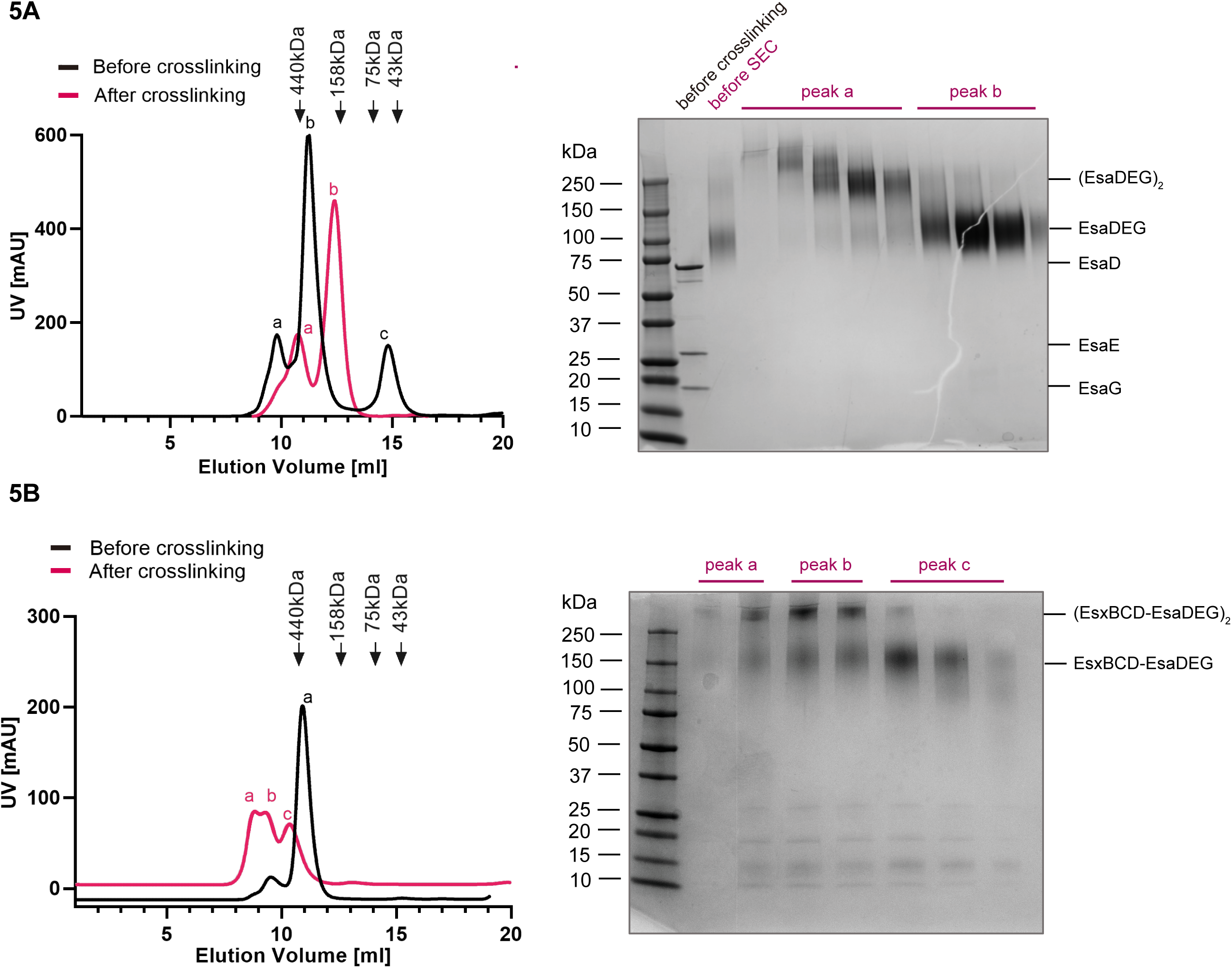
Purification and crosslinking of EsaD complexes for cryoEM. A. Left hand panel: SEC analysis of EsaDEG complexes before (black trace) and after (red trace) glutaraldehyde crosslinking. Right hand panel: SDS PAGE and Coomassie staining of SEC fractions of the crosslinked complexes shows EsaDEG trimers (∼130 kDa) in peak b and (EsaDEG)_2_ hexamers (∼260 kDa) in peak a. B. Left hand panel: SEC analysis of EsaDEG-EsxBCD complexes before (black trace) and after (red trace) glutaraldehyde crosslinking. Right hand panel: SDS PAGE and Coomassie staining of SEC fractions of the crosslinked complexes shows EsaDEG-EsxBCD hexamers (∼150 kDa) in peak c, and a mixture of hexamers and (EsaDEG-EsxBCD)_2_ dodecamers (∼300 kDa) in peaks a and b.

Analysis of crosslinked EsaDEG trimer by single particle cryo-EM yielded reference-free 2D class averages that lacked secondary structure detail but yielded a bilobed shape of approximately 100 Å in diameter. Generation of 3D classes *ab initio* from these particles gave volumes consistent in shape and particle number (21.4 – 29.2 % across four classes), suggesting considerable conformational heterogeneity within the sample (Fig. S3A). 3D reconstructions of the two most populated classes generated low resolution volumes consistent with the 2D class averages (Fig. S3A). An Alphafold model of EsaDEG (Fig. S3C) could be partially docked into one of these volumes (Fig. 6A). In this model EsaE is sandwiched between the N-(residues 1-388) and C-terminal (residues 444-614) domains of EsaD and EsaG interacts almost exclusively with an orthogonal face on the C-terminal domain of EsaD. A recent study showed that a complex between the EsaD nuclease domain and EsaG adopts an alternative conformation to that generated by Alphafold, where the nuclease domain is split into two parts with EsaG sandwiched between them.^26^ However, the overall volume of this domain is similar in both conformations.

**Figure 6.**
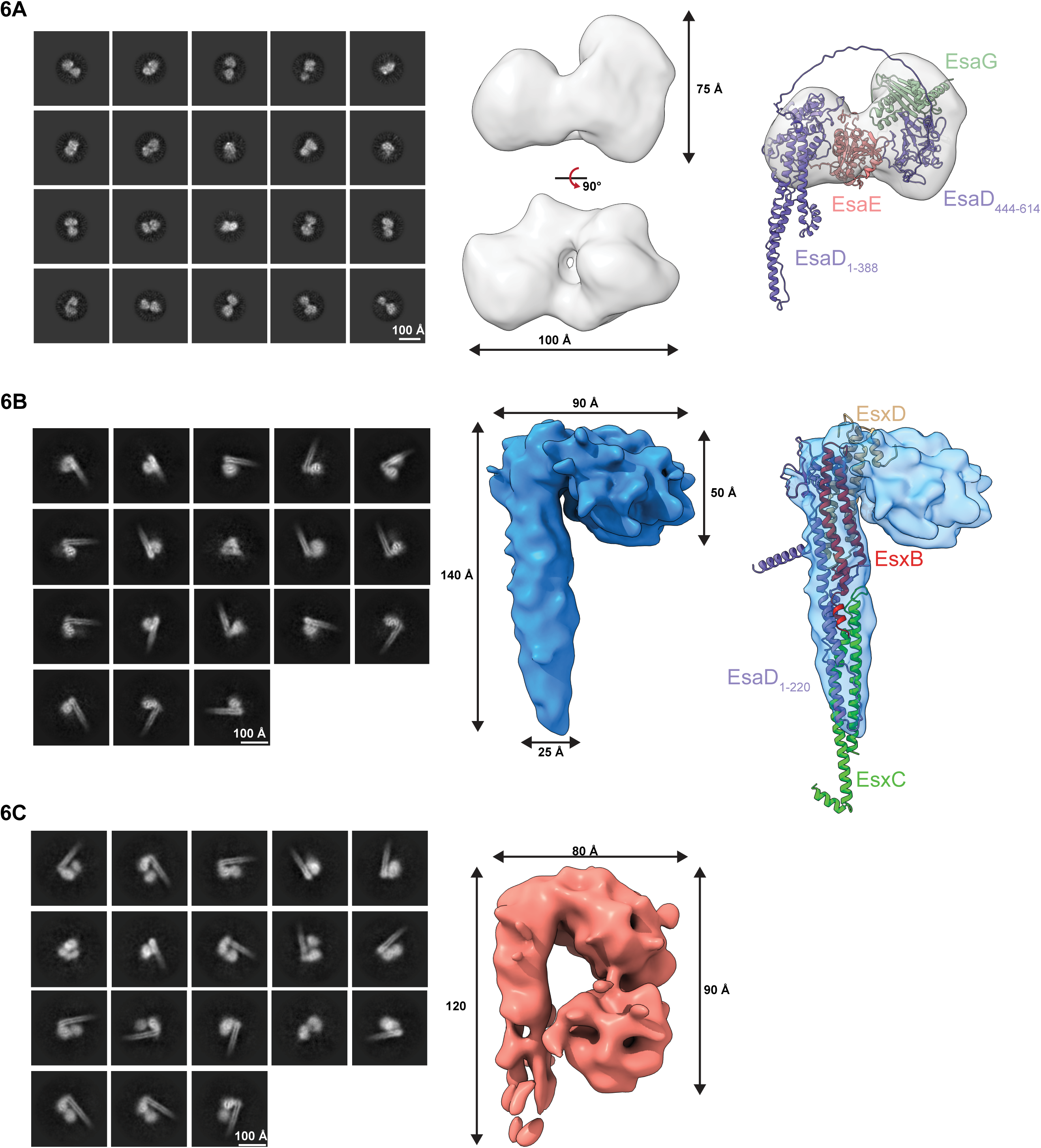
Structural analysis of EsaD complexes by cryo-EM. A. Left, 2D class averages of crosslinked EsaDEG. Middle, low resolution cryo-EM reconstruction of EsaDEG (grey). Right, Alphafold model of EsaDEG partially fit into the cryo-EM volume. Cryo-EM map of EsaDEG shown in transparent grey, with overlayed ribbon models of EsaD (purple), EsaE (pink) and EsaG (green). B. Left, 2D class averages of crosslinked EsaDEG-EsxBCD. Middle, low resolution cryo-EM reconstruction of EsaDEG-EsxBCD (blue). Right, alphafold model of EsaD_1-220_-EsxBCD fit into the cryo-EM volume. Cryo-EM map of EsaDEG-EsxBCD shown in transparent blue, with overlayed ribbon models of EsaD_1-220_ (purple), EsxB (red), EsxC (lime green) and EsxD (beige). C. Left, 2D class averages of an alternate volume of crosslinked EsaDEG-EsxBCD. Right, low resolution cryo-EM reconstruction of alternate EsaDEG-EsxBCD volume shown in light pink.

Single particle cryo-EM was also performed on crosslinked EsaDEG-EsxBCD hexamer. Reference-free 2D class averages included cane-like structures consisting of a helical elongated shaft and lobed head not previously seen with the crosslinked EsaDEG trimer sample (Fig. S3B). One of the four volumes generated *ab initio* from these particles was consistent with the bilobed volume generated from the EsaDEG trimer sample (Fig. S3B, grey volume). The other three *ab initio* classes yielded volumes consistent with cane-like structures, but the size and morphology of the lobed head differed across the classes (Fig. S3B). Refinement of the most populated class yielded a low-resolution volume consisting of a 25 Å x 140 Å shaft and 50 Å by 90 Å lobed head (Fig. 6A). An Alphafold model of EsaD_1-220_EsxBCD matched the dimensions of the shaft and could be docked into this map (Fig 6B); we therefore propose that the shaft of the cane is assembled from these proteins. An alternate volume generated from the second most populated class showed additional density within the cane head (Fig. 5E) that swings back down to contact the shaft. This bilobed domain is grossly consistent in shape and size with the EsaDEG trimer volume. We propose that these two prominent volumes represent separate states of the EsaDEG-EsxBCD hexamer where the toxin-immunity domain is either tethered to the shaft (latter volume) or are partially disordered or disassembled (former volume). These experiments provide the first structural information for a complete pre-secretion complex for a T7b toxin substrate.

### EsaE is structurally homologous to the T7SSa chaperone EspG

In the absence of high-resolution structural data, we examined Alphafold models of individual components of the pre-secretion complex. EsxB, EsxC and EsxD all fold as α-helical hairpins (Fig. S1). For EsaD on its own (Fig S1) or in complex with EsaE and EsaG (Fig S3C) only the nuclease domain could be modelled with high confidence. However, the N-terminal region (residues 1-220) could be confidently modelled in complex with EsxB, EsxC and EsxD, generating a structure that fits well with the cryo-EM determined volume of the shaft (discussed above) (Fig. S3C, S4 and 6B). In this model EsaD_1-220_ extends along the length of the shaft. EsxC binds to the first half of the EsaD helical domain at the bottom end of the shaft and EsxB and EsxD are positioned at the top, adjacent to the globular head. EsxB and EsxD, which have been shown experimentally to interact^20^, form an antiparallel heterodimer within the complex, with the same architecture as known WXG100 dimer structures.^27, 28^ The central WXG motif of EsxB is located at the top end of the complex, and the C-terminus of EsxD, carrying the putative targeting signal YxxxE, protrudes from the shaft at the same end (Fig. S4). This dimer interacts with the second half of the EsaD helical domain, with the widest part comprising a bundle of five α-helices: two each from EsxB and EsxD, and one from EsaD. While the structure is largely α-helical, midway down the shaft are two short antiparallel β-strands contributed by the N-termini of EsxB and EsaD. Each is four residues long, and they are sandwiched between the central loop regions of EsxC and EsxD.

Intriguingly, a search of the PDB using the Dali server revealed the Alphafold model for EsaE to be structurally homologous to EspG chaperones utilized by PPE substrates of the mycobacterial T7SSa^29–31^, despite lacking any sequence homology to these proteins. The top three hits from the Dali search have Z-scores of 10.3 (EspG3, PDB 5SXL), 10.0 (EspG5, PDB 4W4L) and 9.9 (EspG1, PDB 5VBA), and Figure S1C shows a structural alignment of the EsaE model with the EspG3 crystal structure, with an RMSD of 3.2 Å. EspG proteins are found in mycobacterial T7a systems and, like EsaE, function as targeting chaperones for T7 substrates. EspG substrates belong to the PE/PPE protein families which, like LXG proteins, have a conserved N-terminal domain and a highly polymorphic C-terminal cargo domain.^32^ In the published crystal structure of a trimer of PE25, PPE41 and EspG3 from *M. tuberculosis*, the PE/PPE dimer is a long narrow bundle of four α-helices to which EspG3 binds at one end^30^ and bears a striking resemblance to our cryo-EM structure (Fig S4). Binding of EsaE to EsaD might therefore mimic the interaction between EspG and the PE/PPE substrates of the T7SSa.

## Discussion

Previously it has been shown that the EsaD nuclease toxin forms a ternary complex with a chaperone, EsaE, and an immunity protein, EsaG.^9^ Here we show that EsaD has three additional protein partners, EsxB, EsxC and EsxD that interact with the EsaD N-terminal region, facilitating its secretion. Cryo-EM analysis of the EsaDEG and EsaDEG-EsxBCD complexes indicate that incorporation of the small partner proteins results in the formation of a complex with a long helical shaft and a globular head. The shaft has a diameter of approximately 25 - 30 Å, similar to the diameter of the EsxA homodimer that is known to be secreted in a folded form.^28, 33^ A model for EsaD secretion is shown in Fig. 7. It should be noted that current high-resolution structures of the hexameric T7SSa show a closed central pore.^16, 17^ While the size of the open channel is not clear, modelling studies, based at least in part on high resolution structures of the related hexameric DNA translocase FtsK, suggest it may be in the region of 30 Å.^34^ We propose that the helical shaft is secreted in a folded conformation (Fig. 7), which would account for the co-dependent secretion of these proteins observed previously.^20^ Intriguingly, our Cryo-EM analysis revealed that the EsaD cargo domain (comprising the EsaD nuclease domain, EsaG and EsaE) exists in at least two different states, one of which is partially disordered or disassembled (Fig. 6BC, Fig. S3B), supporting the view that this domain is not tightly folded. As the dimensions of the cargo domain are significantly wider than the predicted pore size, there would likely need to be some unfolding of this part of the complex to be accommodated within the secretion channel. The EsaG immunity protein may be liberated from the complex at this point, as it is not secreted with EsaD.^9^

**Figure 7.**
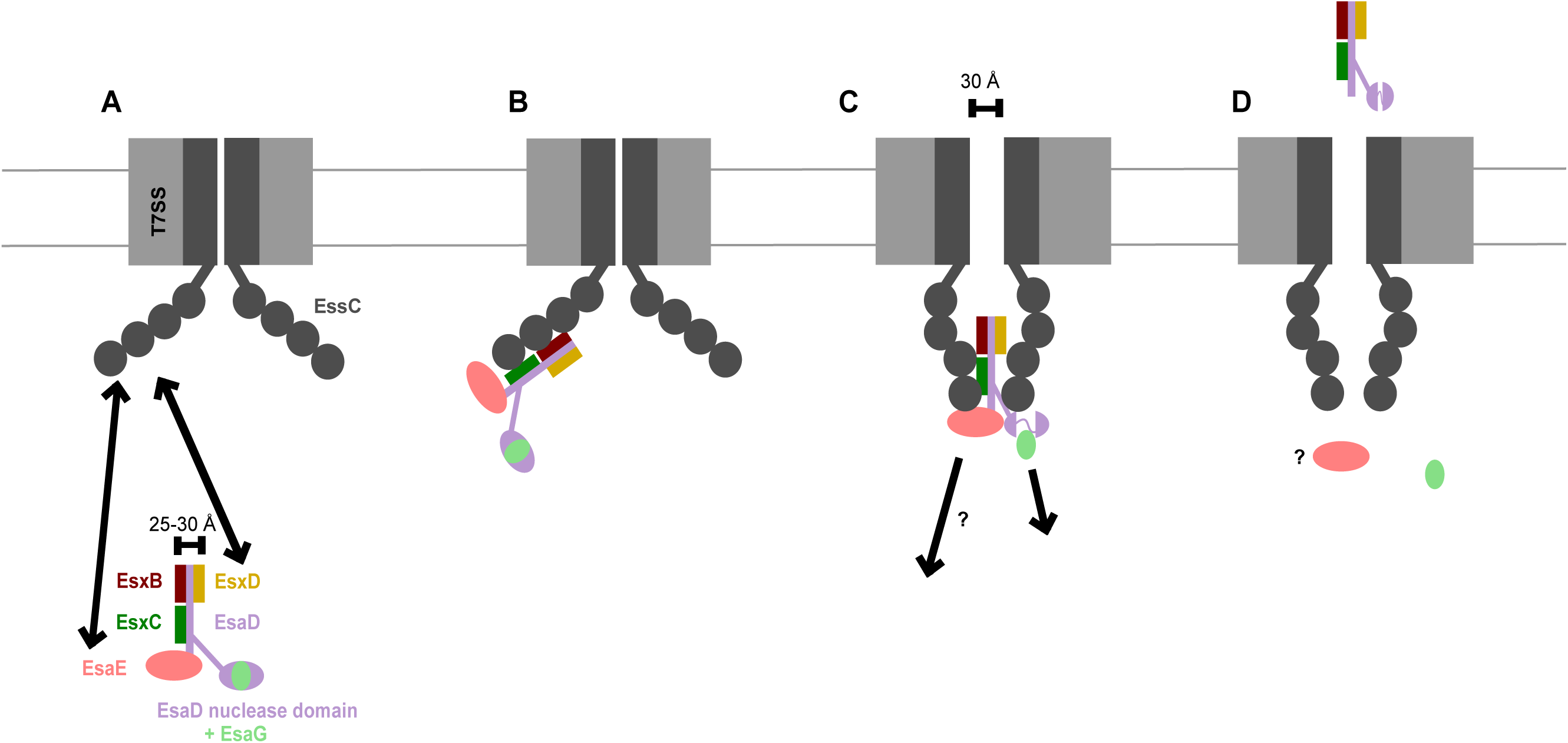
– Model for EsaD secretion by the T7SS. A. The EsaD pre-secretion complex engages the ATPase domains of EssC mediated through interaction of both the chaperone EsaE, and the EsxD C-terminus with EssC. B. Interaction of the pre-secretion complex with the EssC ATPase domains promotes a conformational change to a ‘contracted’ conformation where the ATPase domains from adjacent protomers are brought into close proximity to form a cage.^16^ C. The membrane pore opens to allow passage of the secretion complex. We hypothesize that the cage/pore is too narrow to allow passage of the folded EsaD nuclease domain with bound EsaG and that the nuclease domain unfolds around the active site, as described previously^26^, releasing the immunity protein. EsaE many also be released at this stage. D. Secreted EsaD with its EsxB, EsxC and EsxD partner proteins are released at the surface of the cell, potentially facilitated by the T7SSb component EsaA^37^ (not shown). Note that ATP binding/hydrolysis by EssC and interaction with EsxA are also expected to be required for at some stages the secretion process, but these have been omitted for clarity.

WXG100-like protein partners have previously been identified for T7SSb substrates with N-terminal LXG domains from *Streptococcus intermedius*.^10, 35^ Two small WXG100-like proteins designated Lap1 (a DUF3130 family protein) and Lap2 (a DUF3958 protein) are encoded upstream of each *S. intermedius* LXG toxin, and these function as export factors forming a pre-secretion complex with the LXG region of their cognate toxin.^35^ Lap1 carries a C-terminal FxxxD targeting signal, similar to the YxxxD/E targeting signal in T7a substrates.^22^ While the N-terminal helical domain of EsaD is not recognized as belonging to the IPR006829 LXG domain family, a PFAM LXG domain signature (PF04740) is detected from EsaD residue 1-71, and an HHPred search of the PFAM database predicts a PF04740 domain from residues 2-170 with 98% probability. We therefore conclude that the first 170 residues of EsaD do encompass an LXG domain, but this is more divergent than those categorized as IPR0044927, such as the LXG domains of the *S. intermedius* TelABCD toxins. While there are no DUF3130- or DUF3958-encoding genes at the *esaD* locus, we show here that EsxB, EsxC and EsxD function analogously to Lap1 and Lap2 in EsaD secretion. Furthermore, a C-terminal arm including a YxxxE motif is found in EsxD and likely functions in EsaD targeting to the translocon. The reason for the differing requirement for two or three WXG100-like proteins in a pre-secretion complex is not clear, although this may correlate with structural differences between different types of LXG domain.

The T7SSa system, found in actinobacteria, and T7SSb system present in firmicutes share limited similarity at the sequence level. Both utilize an FtsK/SpoIIIE family ATPase for substrate recognition and protein transport, albeit the T7SSb EssC carries two N-terminal forkhead-associated (FHA) domains that are absent from the T7SSa homologue EccC. Additionally, both systems encode at least one WXG100 family protein, although the sequence similarity between these components is low. Beyond EssC/EccC and EsxA/ESAT-6, the two systems utilize a number of additional core components which are completely unrelated. We have shown here, from a combination of cryo-EM and Alphafold modelling, that the structure of the T7SSb EsaD pre-secretion complex shares a common architecture with the PE-PPE-EspG pre-secretion complexes of mycobacterial T7SSa systems, despite a complete lack of sequence similarity. This includes a long α-helical shaft with a diameter of ∼25 Å. A long α-helical shaft has also been seen in the crystal structure of the T7SSa substrate EspB.^36^ This striking structural similarity suggests an unexpected level of mechanistic overlap, which will contribute to our understanding of both systems. Within each type of complex, targeting information is supplied by both a WXG100-like partner and an EsaE/EspG partner. These targeting signals presumably interact with the EssC ATPase domains which are conserved in both systems. It will be interesting to see whether the structural features we describe here are common to other substrates of the T7a and T7b systems.

## Acknowledgements

This study was supported by the Wellcome Trust (through Investigator Awards 10183/Z/15/Z and 224151/Z/21/Z to TP), the Intramural Research Program of the NIH, NCI, Center for Cancer Research (awarded to SL), the China Scholarship Council (through award of a PhD studentship to YY) and the University of Newcastle. We thank the bioscience technology facility at the University of York for mass spectrometry and analysis.

## Author contributions

Conceptualization (YY, EB, FA, TP); Methodology (YY, EB, JD, FA, SL, TP); Investigation (YY, EB, JD); Writing – original draft (YY, EB, JD, FA, SL, TP) and funding acquisition (TP and SL).

## Materials and Methods

### Bacterial strains, plasmids and growth conditions

*E. coli* was cultured in LB medium (Melford) with ampicillin (110 ug/ml) where required; S. aureus was cultured in tryptic soy broth (TSB) (Oxoid) with chloramphenicol (10 ug/ml) where required.

All strains, plasmids and primers used in this study are listed in Tables S1, S2 and S3 respectively. Plasmids for expression in *S. aureus* for secretion assays were based on pRab11^38^ and were constructed as follows: *esaG* was amplified from COL genomic DNA and cloned into pRab11 KpnI and EcoRI restriction sites to make pRab11-esaG. *esaD-HA* was then amplified and inserted into upstream of *esaG* using HiFi DNA assembly (NEB). The catalytic residue H528 of *esaD* was mutated to encode an alanine using a Q5 site-directed mutagenesis kit (NEB). *esxCBED* amplified from COL genomic DNA was then inserted upstream of *esaD* by HiFi assembly, creating pRab11-CBED-DG. Individual genes were then deleted from this construct using a Q5 site-directed mutagenesis kit.

Plasmids used for protein purification were based on pQE70 (Qiagen). For purification of EsaDEG, genes for SAOUHSC_00267 (*esaE*), SAOUHSC_00268 (*esaD*) and SAOUHSC_00269 (*esaG*) were ordered as gblocks (IDT) codon optimized for *E. coli* expression with a strong RBS attached to each gene. PCR fragments were assembled by HiFi DNA assembly to include two copies of *esaG* followed by one copy of *esaE* and one copy of *esaD* with a C-terminal hexahistidine tag and the H528A substitution, generating pQE70-GGED. The *esaD* gene was amplified from the EsaD gblock using the combinations of codoptDnostop_fwd and codoptH528A_rev or codoptH528A_fwd and codoptDnostop_rev to create fragments that were then joined together by short overlap extension PCR to introduce the H528A mutation using the outside codoptDnostop_fwd and codoptDnostop_rev primers. To make pQE70-GGED-his-CBED-tstrep a genomic fragment encompassing SAOUHSC_00264 (*esxC*), SAOUHSC_00265 (*esxB*), SAOUHSC_00266 (*esaE*) and SAOUHSC_00267 (*esxD*), including the *esxC* RBS, was amplified from *S. aureus* COL genomic DNA. A twin-strep tag was included at the EsxD C-terminus to allow for an additional affinity chromatography step and immunodetection. This fragment was inserted downstream of *esaD* in pQE70-GGED by HiFi assembly. To avoid EsaD toxicity during cloning, all cloning steps were performed in DH5α carrying plasmid pREP4 (Qiagen) for expression of the Lac repressor. The completed plasmids were transformed into *E. coli* M15 pREP4 for protein expression.

Chromosomal deletion of *esxC-esaG* was accomplished using plasmid pIMAY-Z and following the published protocol.^39^ pIMAY-Z_CBEDDG, carrying regions flanking *esxC-esaG*, was constructed by HiFi assembly with primers listed in Table S3. The COLΔ*esxC-esxG* genotype was verified by sequencing of the relevant region.

### Cell fractionation and Western blotting

To monitor EsaD secretion, pRab11-based plasmids encoding EsaD with the catalytic inactivating H528A substitution and a C-terminal HA tag for immunodetection, alongside different combinations of EsxBCD, EsaE and EsaG were transformed into *S. aureus* strain COLΔ*esxC-esxG* by electroporation. Strains were subcultured at 1/25 from an overnight-grown preculture into fresh TSB medium. At OD600nm = 2, cells were collected and the supernatant samples precipitated with trichloroacetic acid in the presence of deoxycholate, as described previously.^24^ Samples were normalized to OD600 = 2 in PBS, and lysed by the addition of 50 μg ml–1 lysostaphin (Ambi) with incubation at 37 °C for 30 min. All samples were mixed with an equal volume of NuPAGE LDS sample buffer and boiled for 10 min. Antibodies against EsxA, TrxA, EsxB, EsxC used for immunoblotting have been previously described.^25, 40, 41^ Antibodies against the HA tag, strep tag and myc tag were obtained from Sigma, IBA and Abcam, respectively.

### Protein purification

Protein complexes were produced from the relevant pQE70 plasmid (see Table S2) in strain M15 pREP4 (Qiagen) as follows: Cells were cultured in LB broth supplemented with ampicillin (125 ug/ml) and kanamycin (50 ug/ml) at 37°C with aeration until an OD_600nm_ of 0.6-1.0 was reached, before induction with 0.5 mM IPTG and growth overnight at 18°C. Cells were harvested by centrifugation and lysed by sonication in binding buffer (50 mM HEPES pH 7.5, 150 mM NaCl and 50 mM imidazole) with cOmplete protease inhibitor tablets (Roche). Cell lysates were clarified by centrifugation and applied to a 5ml Histrap FF column (Cytiva) equilibriated in the same buffer. The column was washed with binding buffer and bound proteins were eluted using a 50-500 mM imidazole gradient. Eluted protein was concentrated using a 10 kDa MWCO spin concentrator (Vivaspin) before injection onto a Superdex S200 10/300 gl column equilibrated in 50 mM HEPES pH 7.5, 150 mM NaCl. The hexameric complex containing full length EsaD carried an additional twin-strep tag on EsxD, and this complex was further purified by strep-tactin affinity purification, followed by a second round of SEC.

For glutaraldehyde crosslinking the SEC fractions corresponding to major peaks were diluted to 0.5 mg/ml in 50 mM HEPES pH 7.5, 150 mM NaCl and crosslinked using 0.1% glutaraldehyde for 30 minutes at room temperature, before quenching with at least a tenfold excess of tris. The crosslinked protein was concentrated again using a 10 kDa MWCO spin concentrator before injection on Superdex S200 10/300 gl. Eluted protein was again concentrated by spin concentrator (10 kDa MWCO) and snap frozen with liquid nitrogen for subsequent cryo-EM analysis.

### Cryo-EM sample preparation and data acquisition

Crosslinked EsaDEG and EsaDEG-EsxBCD samples were diluted to 0.05 mg/ml and 0.2 mg/ml in SEC buffer, respectively. Three microliters of sample were applied onto glow-discharged (60 s, 30 mA) 300-mesh R1.2/1.3 Quantifoil Au grid. Grids were blotted for 2 s in 100 % humidity at 8-12 ℃ and plunge frozen in liquid ethane using a Vitrobot Mark IV (Thermo Fisher Scientific). Data were collected in counted mode in EER format on a CFEG-equipped Titan Krios G4 (Thermo Fisher Scientific) operating at 300 kV with a Selectris imaging filter (Thermo Fisher Scientific) with slit width of 10 e^-^V and Falcon 4 direct detection camera (Thermo Fisher Scientific) at ×165,000 magnification, with a physical pixel size of 0.723 Å (EsaDEG) or 0.693 Å (EsaDEG-EsxBCD). Movies were collected with a total dose of 57.0 e^-^/Å^2^.

### Cryo-EM data processing

Patched motion correction, CTF parameter estimation, particle picking, extraction, and initial 2D classification were performed in SIMPLE 3.0.^42^ All downstream processing was carried out in cryoSPARC v3.3.1^43^, including Gold-standard Fourier shell correlations (FSCs) using the 0.143 criterion. For crosslinked EsaDEG trimer (Fig. S3A), 3,493,119 particles were extracted from 5,824 motion-corrected micrographs and subjected to reference-free 2D classification (k=500). An additional round of reference-free 2D classification was performed on 678,695 pruned particles in cryoSPARC (k=200). 2D-cleaned particles (278,959) were then used to generate four volumes *ab initio*. Particles belonging to the two most populated classes were individually selected for non-uniform refinement against their 30 Å lowpass-filtered volume counterparts, yielding volumes of moderate resolution from which only gross conformational information could be assessed. For crosslinked EsaDEG-EsxBCD hexamer (Fig. S3B, 2,615,890 particles were isolated after an initial round of reference-free 2D classification (k=300). These particles were further cleaned with another two rounds of reference-free 2D classification within cryoSPARC (k=200 each round). Particles (343,131) belonging to classes with strong structural features were selected and used to generate four volumes *ab initio*, of which one class was similar to the volume generated from the crosslinked EsaDEG trimer. The other three classes demonstrated cane-like structures that consisted of a shaft and head. The head varied in size across the three classes, and the two most well-defined classes were further selected for non-uniform refinement against their own 15 Å lowpass-filtered volumes as references. Refinements produced moderate resolution volumes of cane-like structures consisting of a shaft and either a large or bilobed head. Further extensive classification in 2D and 3D space for both the EsaDEG trimer and EsaDEG-EsxBCD hexamer datasets did not significantly improve map quality, indicating the samples suffered from conformational and/or compositional heterogeneity.

### Modelling and map fitting

Alphafold was used to generate models of individual proteins.^44^ Alphafold models of EsaDEG and EsaD_1-220_EsxBCD (Fig. S3C) were generated using the default settings of Alphafold-Multimer^45^ using UniProt entries Q2G179 (EsaD), Q2G181 (EsaE), Q2G179 (EsaG), Q2G182 (EsxB), Q2G183 (EsxC), Q2G180 (EsxD). The top ranked Alphafold models were fit into the cryo-EM maps using the fitmap function of USCF ChimeraX (43). The superose program within ccp4 was used to align EsaE with EspG3. Map and model figures were prepared within ChimeraX.^46^

### Flanking gene analysis (FlaGs)

Gene neighborhood analysis was carried out using the WebFlaGs server.^47^ Input sequences were generated by running a BLASTP search on bacilli, excluding *Staphylococcus*, with *S. aureus* EsaD (Q2G179) as a query. Sequences lacking the LXG domain (residues 2-120) were manually removed, and the resulting sequence list was input into WebFlaGs with ten flanking genes selected and other parameters set to default.

