## Supplemental Figures and Tables for "Three small partner proteins facilitate the type VII-dependent secretion export of an antibacterial nuclease"

Tracy Palmer

**Supporting Information**

**This PDF file includes:**

Figures S1 to S4

Tables S1 to S3

SI References

Fig. S1. Alphafold structures of EsaD and its accessory proteins

A. Alphafold models for each of the indicated proteins from *S. aureus* strain COL.

B. Alphafold model of WP_185417595, the EsxC protein from *Listeria booriae*, shown as an open pink arrow in Figure 1D. Models in (A) and (B) are colored according to the Alphafold model confidence.

C. Structural alignment of EsaE (modelled by Alphafold) (orange) with the crystal structure of EspG3 (PDB 4L4W). The aligned atoms have an RMSD of 3.2 Å.

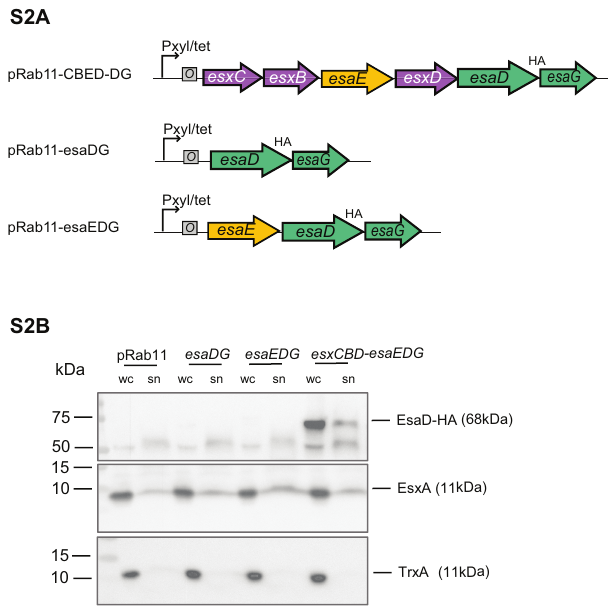

Fig. S2. Contribution of EsxBCD to EsaD secretion.

A. Schematic showing the constructs used in (B).

B. COL *ΔesxC-esaG* cells carrying empty vector (pRab11), or pRab11 carrying the indicated genes, were cultured in TSB growth medium. Following induction of plasmid-encoded gene expression, whole cell (wc) and culture supernatant (sn) fractions were isolated, prepared for immunoblot, and separated by SDS PAGE. EsaD was detected with antibodies against the C-terminal HA tag (upper panel). The T7-secreted control EsxA and cytoplasmic control TrxA were detected with antibodies against EsxA (middle panel) and TrxA (lower panel). N.B. the supernatant fractions analyzed here correspond to 6 x more culture volume than the cell fractions.

Fig. S3. Cryo-EM workflows

S3A. Cryo-EM image processing workflow for EsaDEG

S3B. Cryo-/EM image processing workflow for EsaDEG-EsxBCD

S3C. Alphafold models used for docking, colored by pLDDT score. Left, EsaDEG. Right, EsaD_1-220_-EsxBCD.

Fig. S4. Architecture of T7a and T7b substrate complexes.

An Alphafold model for EsaD_1-220­_-EsxBCD (left) alongside the crystal structure of a trimer of PE25, PPE41 and EspG3 from *M. tuberculosis* (right) (PDB 4W4L).^1^

Table S1. Strains used in this study

| **Strain** | **Genotype/description** | **Source/Reference** |
| --- | --- | --- |
| COL | *essC1* *S. aureus* strain; MRSA *agr* | Reference^2^ |
| COL Δ*esxC-esxG* | COL Δ*SACOL0277 – SACOL0282* | This study |
| M15 (pREP4) | Commercial *E. coli* strain for expression from pQE plasmids. *F-, lac, ara, gal, mtl* [*KanR, lacI*] | Qiagen |
| JM110 | *rpsL* (Strr) *thr leu thi-1 lacY galK galT ara tonA tsx dam dcm supE44* ∆(*lac-proAB*) [F ́ *traD36 proAB lacI*q*Z*∆*M15*] | Lab stock |
| DH5α | *Δ(argF-lac)169, φ80dlacZ58(M15), ΔphoA8, glnX44(AS), deoR481, rfbC1, gyrA96(NalR), recA1, endA1, thiE1 and hsdR17* | Lab stock |

Table S2. Plasmids used in this study

| **Plasmid** | **Genotype/description** | **Source/Reference** |
| --- | --- | --- |
| pRab11 | *E. coli* / *S. aureus* shuttle vector, P_xyl/tet_ inducible promoter, cml^r^, amp^r^ | Reference^3^ |
| pRab11-CBED-DG | pRab11 carrying *esxC*, *esxB*, *esaE*, *esxD, esaD^H528A^-HA*, *esaG* for tetracycline-inducible expression | This work |
| pRab11-CED-DG | pRab11-CBED-DG with *esxB* deleted | This work |
| pRab11-BED-DG | pRab11-CBED-DG with *esxC* deleted | This work |
| pRab11-CBE-DG | pRab11-CBED-DG with *esxD* deleted | This work |
| pRab11-CBD-DG | pRab11-CBED-DG with *esaE* deleted | This work |
| pRab11-esaDG | pRab11 carrying*, esaD^H528A^-HA*, *esaG* for tetracycline-inducible expression | This work |
| pRab11-esaEDG | pRab11 carrying *esaE*, *esaD^H528A^-HA*, *esaG* for tetracycline-inducible expression | This work |
| pIMAY-Z | *E. coli* / *S. aureus* shuttle vector, temperature sensitive, cml^r^ | Reference^4^ |
| pIMAY_esxC-esaG | pIMAY-Z carrying *SACOL0277 – SACOL0282* deletion allele | This work |
| pQE70-GGED | pQE70 encoding two copies of SAOUHSC_00269 (EsaG), one each of SAOUHSC_00266 (EsaE) and SAOUHSC_00268 (EsaD) with a H528A mutation and a hexahistidine C-terminal tag. All genes codon optimised for *E. coli* expression. | This work |
| pREP4 | *lacI KanR* | Qiagen |
| pQE70-GGED-his-CBED-tstrep | pQE70 carrying *esxC*, *esxB*, *esaE*, *esxD-^twinstrep^, esaD^H528A^-HA*, *esaG* for IPTG-inducible expression | This work |
| pQE70-GGED(1-477)-his-CBED | pQE70-GGED-his-CBED with EsaD toxin domain deleted | This work |
| pQE70-GGED(478-614)-his-CBED | pQE70-GGED-his-CBED­­­ with EsaD N-terminus deleted | This work |

**Table S3.** Primers used in this study

| **Construct** | **Primer** | **sequence** |
| --- | --- | --- |
| pRab11-esaG | EsaG For | GGATTGGTACCAGGAGGTTTCTAGTT ATGACATTTGAAGAGAAGCTTAGCA |
|  | EsaG Rev | GCTAAGAATTCTTATTCTTCTAGCTCTTTAATATAT |
| pRab11-esaDG | esaG_fwd | AGGAGGTTTCTAGTTATGAC |
|  | esaG_rev | GGTACCATCATACTCTATCAATG |
|  | EsaD_fwd | tgatagagtatgatggtaccaggaggtttctagttATGACAAAAGATATTGAATATCTAAC |
|  | EsaD_rev | gtcataactagaaacctcctctaagcgtaatctggaacatcgtatgggtaCTTATTTAATATTCTTCTAATATTTCTTTCAC |
| pRab11-esaDG | H528 | TGATGGAGGTGCCTTAATCGCTAG |
| (H528A substitution) | H528 | TCGTCTGGTAATCTATCC |
| pQE70-GGED | codoptG1_fwd | ATTCATTAAAGAGGAGAAATTAAGCATGACATTTGAAGAAAAACTTAGC |
|  | codoptG1_rev | ATTCATTAAAGAGGAGAAATTAAGCATGACATTTGAAGAAAAACTTAGC |
|  | codoptG2_fwd | CTGGAAGAGTAATAAATTAAAGAGGAGAAATTAACCATG |
|  | codoptG2_rev | CTCCTCTTTAATGTTACTCTTCCAGCTCCTTAATG |
|  | codoptE_fwd | CTGGAAGAGTAACATTAAAGAGGAGAAATTAAGC |
|  | codoptE_rev | AATGGGCCCTTATTATTCTTCTGCCTTGTTG |
|  | codoptDnostop_fwd | GGCAGAAGAATAATAAGGGCCCATTAAAGAG |
|  | codoptDnostop_rev | AAGCTTAGTGATGGTGATGGTGATGCTTATTCAGAATACGACGG |
|  | pQE70_GGEDco_fwd | CATCACCATCACCATCAC |
|  | pQE70_GGEDco_rev | GCTTAATTTCTCCTCTTTAATG |
|  | codoptH528A_fwd | GACGACGGTGGAGCCCTGATTGCTCGC |
|  | codoptH528A_rev | GCGAGCAATCAGGGCTCCACCGTCGTC |
| pQE70-his-CBED-tstrep | QE backbone_rev | GCTTAATTTCTCCTCTTTAATG |
|  | CBED-fwd | ttaaagaggagaaattaagcatgcatcaccatcaccatcacAATTTTAATGATATTGAAACAATGGTTAAG |
|  | CBED_rev | tggatctatcaacaggagtcctacagatcctcttctgagatgagtttttgttcTCCCTCAATATTATAGTAAAGC |
| pQE70-GGED (1-477)-CBED | 9 delete 478-614_F | CATCACCATCACCATCAC |
|  | 9 delete 478-614_R | TGTATATTCAATATTTGCTTTAAG |
| pQE70-GGED-(421-614)-CBED | 13Delete 1-420_F | ACCCATGGCCCAAAGGAT |
|  | 13Delete 1-420_R | CATGGTTAATTTCTCCTCTTTAATGG |
| pRab11-CBED-DG | 1DG_fwd | AGGAGGTTTCTAGTTATGAC |
|  | 1DG_rev | GTACCATCATACTCTATCAATG |
|  | CBED_F | ttgatagagtatgatggtacATGAATTTTAATGATATTGAAACAATG |
|  | 2CBED_R | gtcataactagaaacctcctCTATCCCTCAATATTATAGTAAAGC |
| pRab11-CED-DG | EsxB-deleted | tctagttATGAAAGATGTTAAGCGAATAG |
|  | EsxB-deleted _R | aacctcctTTAATTCATTGCTTTATTAAAATATTC |
| pRab11-BED-DG | EsxC-deleted-F | ATGGGTGGATATAAAGGTATTAAAG |
|  | EsxC-deleted-R | AACTAGAAACCTCCTGTAC |
| pRab11-CBE-DG | EsxD-4th-F | AGGAGGTTTCTAGTTATG |
|  | EsxD-4th-R | TTACTCCTCTGCTTTATTAATATG |
| pRab11-CBD-DG | EsaE-deleted_F | tctagttATGACGTTGAGTGGAAAAATTAG |
|  | EsaE-deleted_R | aacctcctTCATGGGTTCACCCTATC |
| pRab11-esaEDG | nnEDG_fwd | AGGAGGTTTCTAGTTATGAC |
|  | nnEDG_rev | GTACCATCATACTCTATCAATG |
|  | nnE_fwd | ttgatagagtatgatggtacaggaggtttctagttATGAAAGATGTTAAGCGAATAG |
|  | nnE_Rev | gtcataactagaaacctcctTTACTCCTCTGCTTTATTAATATG |
| pIMAY-Z_CBEDDG | pIMAY-Z_fwd | CCGGGGGATCCACTAGTTC |
|  | pIMAY-Z_rev | TCGATATCAAGCTTATCGATACCG |
|  | up-700_fwd | atcgataagcttgatatcgaAGATGAAATTAAATATGAAGATTACAGAG |
|  | up-700_rev | ttaagatagtAACATACCTCCCTCCTATTTAATTC |
|  | down-700_fwd | gaggtatgttACTATCTTAATGTAAGACTAAACAATAAAG |
| ­ | down-700_rev | agaactagtggatcccccggTTTTTCAATATATATAAATGAGCTCAAC |
